# Influence of the microenvironment on the modulation of the host response by the typhoid toxin

**DOI:** 10.1101/2020.06.04.133686

**Authors:** Océane C.B. Martin, Deborah Butter, Eleni Paparouna, Sofia D.P. Theodorou, Maria M. Haykal, Elisa Boutet-Robinet, Toma Tebaldi, Anna Bergonzini, Maria Lopez Chiloeches, Andrew Wakeham, Mikael Rhen, Vassilis G. Gorgoulis, Tak Mak, Ioannis S. Pateras, Teresa Frisan

**Author notes:** Corresponding author Teresa Frisan. These authors shared senior authorship (Ioannis S. Pateras, Teresa Frisan). Univ. Bordeaux, INSERM, UMR1053 Bordeaux Research in Translational Oncology, BaRITOn, Bordeaux, France. TUM School of Life Sciences, Technical University of Munich, Freising, Germany.

## Abstract

Bacterial genotoxins cause DNA damage in eukaryotic cells, resulting in activation of the DNA damage response (DDR) *in vitro*. These toxins are produced by Gram negative bacteria, enriched in the microbiota of Inflammatory Bowel Disease (IBD) and colorectal cancer (CRC) patients. However, their role in infection remains poorly characterized. We have addressed the role of the typhoid toxin in the modulation of the host-microbial interaction in health and disease.

Infection with a genotoxigenic *Salmonella* protected mice from intestinal inflammation. The toxin-induced DNA damage caused senescence *in vivo*, which was uncoupled from the inflammatory response, and associated with the maintenance of an anti-inflammatory environment. This effect was lost when infection occurred in mice suffering from inflammatory conditions that mimic Ulcerative Colitis, a form of IBD.

These data highlight a complex context-dependent crosstalk between bacterial genotoxins-induced DDR and the host immune response, underlining an unexpected role for bacterial genotoxins.

## Introduction

The typhoid toxin (TT) and the cytolethal distending toxins (CDTs), produced by several Gram negative, belong to a family of bacterial protein effectors that cause DNA damage in eukaryotic cells (Grasso and Frisan, 2015). CDTs and TT are tripartite toxins that share the CdtB active subunit, a structural and functional homologue to the mammalian DNase I (Nesic et al., 2004; Song et al., 2013). Upon delivery into the nucleus of the target cells, the CdtB subunit induces DNA strand breaks, which activate the DNA damage response (DDR), coordinated by the sensor kinase Ataxia Telengectasia mutated (ATM) (Grasso and Frisan, 2015). As a consequence of DDR activation, the intoxicated cells are arrested in the G1 and/or G2 phases of the cell cycle and initiate DNA repair. However, if the extent of the damage is beyond repair, most cell types undergo senescence (Blazkova et al., 2010; Ibler et al., 2019; Secher et al., 2013). Occasionally, intoxicated cells may survive and overcome the DDR-induced tumorigenic barrier, leading to genomic instability and acquisition of carcinogenic traits (Guidi et al., 2013; Hanahan and Weinberg, 2011).

DNA damage-induced cellular senescence is characterized by: i) a generally permanent cell cycle arrest, induced by activation of the cyclin-dependent kinase inhibitors p16, p15 and p21, and ii) an active metabolic state resulting in secretion of a broad panel of mediators, most of them with a pro-inflammatory profile, including cytokines (IL1α, IL6, IL8), growth factors (HGF, GM-CSF), and metalloproteases (MMP1-1 and MMP-3) (Gorgoulis et al., 2019). The latter has also been defined as senescence-associated secretory phenotype (SASP) (Campisi and d’Adda di Fagagna, 2007; Krtolica et al., 2001). The pro-inflammatory response induced by DNA damage can be further fuelled by type I IFN secretion induced by the recognition of cytosolic DNA fragments by the cGAS-STING pathway (Hartlova et al., 2015; Li and Chen, 2018).

Considering the mode of action and the plethora of non-cell autonomous effects that senescent cells exert in the surrounding microenvironment, it is puzzling that certain bacteria have acquired genotoxins, since it is unlikely that their primary purpose is to induce/promote cancer in the mammalian host. To understand the role of these bacterial effectors in infection, we have assessed the role of a functional typhoid toxin, produced by *Salmonella* Typhimurium in an *in vivo* model of acute infection.

We demonstrated that infection with a strain expressing a functional typhoid toxin, even though inducing DNA damage, protected mice from intestinal inflammation. The response was characterized by induction of senescence, NFκB activation, reduced recruitment of leukocytes and T lymphocytes, production of Th2 cytokines, higher percentage of anti-inflammatory macrophages and regulatory T cells. This anti-inflammatory effect, partially dependent on a functional ATM, was lost when infection with the genotoxigenic strain occurred in subjects with pre-existing colitis.

These data can have a broader implication since other bacterial genotoxins may also modulate the microenvironment, and the outcome can be different, depending on the bacterium life-style and the tissue tropism.

## Results

### Infection with genotoxin-producing bacteria induces senescence *in vivo*

We have previously demonstrated that infection with a *S.* Typhimurium expressing a functional typhoid toxin enhances host survival and reduces the extent of the colonic inflammation in healthy mice (Del Bel Belluz et al., 2016). We have extended here this analysis by performing a more detailed characterization of the host response in Sv129 mice infected with the genotoxigenic *Salmonella* strain (here defined as MC1 TT) for 5 days and 10 days. As control, we have used an isogenic strain lacking the genotoxigenic CdtB subunit (here defined as MC1 Δ*cdtB*). Presence of a function typhoid toxin enhanced the survival rate of the infected mice compared to mice infected with the control strain (Figure S1A). Consistent with these data, histological evaluation revealed decreased severity of enteritis and colitis along with reduced number of Payer patches in MC1 TT-treated mice, which was observed 10 days post-infection (Figures 1A, 1B and 1C). This effect was not dependent on a different efficiency of colonization of the MC1 TT compared to the control strain (Figure S1B). The reduced inflammatory response was also detected in the small intestine of C57BL/6 mice infected with the attenuated genotoxigenic *Salmonella* strain MC71 (MC71 TT) carrying a mutation of polynucleotide phosphorylase (PNPase) gene (Clements et al., 2002) (Figure S2). Since we did not observe colonic inflammation in this mouse strain, the analysis of the C57BL/6 model was focused on the small intestine.

**Figure 1.**
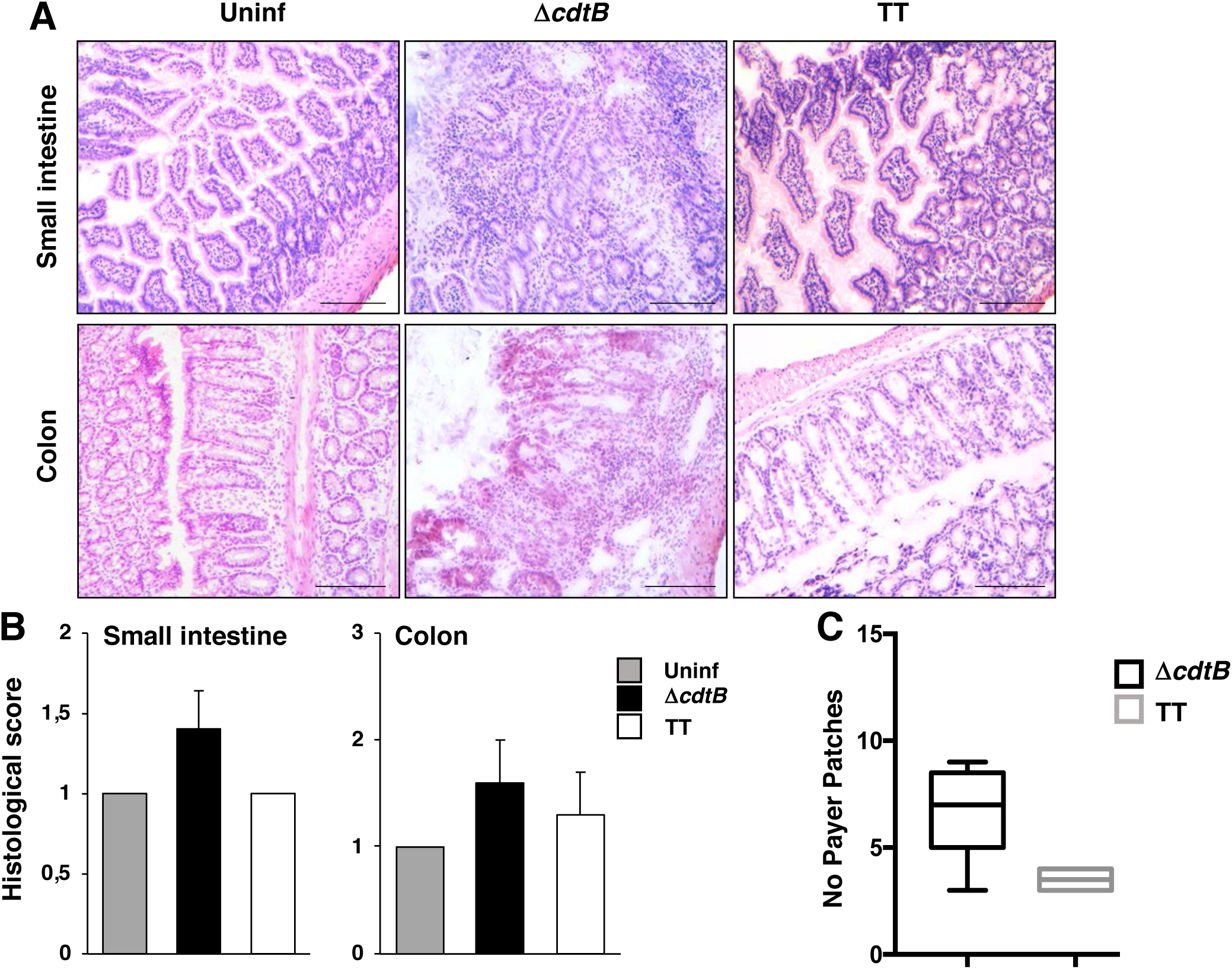
Infection with the typhoid toxin-producing *Salmonella* reduces the inflammatory response. Sv129 mice were mock infected with PBS (Uninf) or infected with the MC1 Δ*cdtB* (Δ*cdtB*) or MC1 TT (TT) strains for 10 days. **A.** Haematoxylin and eosin staining of the small intestine and colon. Scale bar: 200μM. **B.** Histological scores of small intestine and colon, (n mice =5). **C.** Quantification of the macroscopically visible inflamed Peyer Patches, (n mice =5).

These data indicate that the typhoid toxin effect is very robust and can be observed independently on the mouse and *Salmonella* strains.

DNA damage induced by bacterial genotoxins *in vivo* has been commonly evaluated using surrogate markers of the DDR, such as the phosphorylated form of the Histone H2AX (γH2AX) (Rogakou et al., 1999). We have now assessed direct induction of DNA fragmentation in cells isolated from the intestinal mucosa, using an alkaline comet-based assay, considered a gold-standard method to detect DNA strand breaks (Pu et al., 2015). Our data showed that a functional typhoid toxin promotes DNA fragmentation to a greater extent than that observed in mice infected with the control *Salmonella* (Figure 2A). In spite of this observation, we detected enhanced levels of H2AX phosphorylation in the mucosa of mice infected with the MC1 Δ*cdtB* strain (Figure 2B and S3).

**Figure 2:**
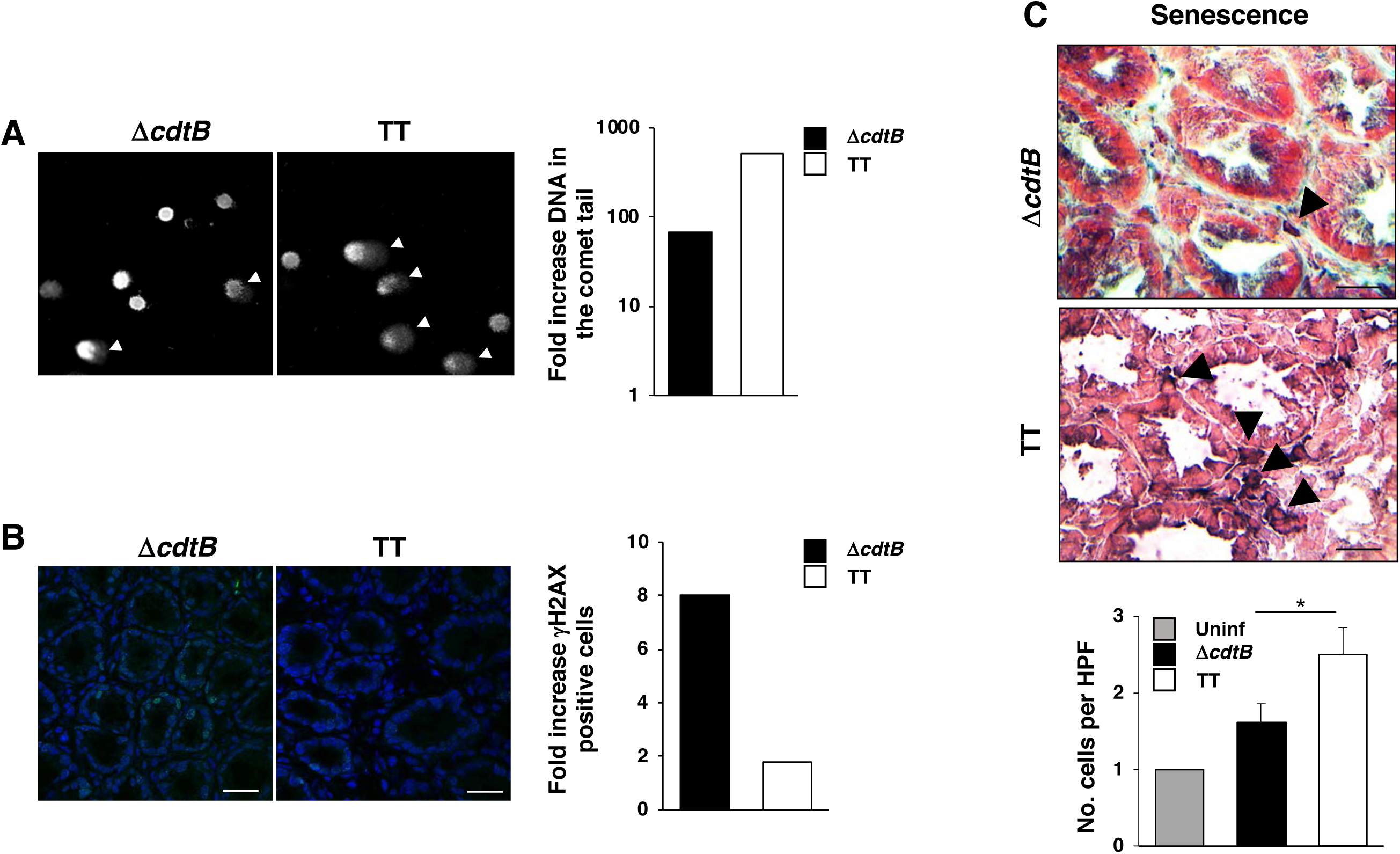
Senescence induced by infection with the genotoxigenic *Salmonella* is uncoupled from the inflammatory response. Sv129 mice were mock infected with PBS or infected with the MC1 Δ*cdtB* (Δ*cdtB*) or MC1 TT (TT) strains for 10 days. **A.** DNA fragmentation. Left panel: representative micrograph of DAPI stained nuclei. Left panel: quantification comet tail: data are reported as ratio between the median value of the % DNA in the comet tail in cells isolated from mice infected with the MC1 Δ*cdtB* or MC1 TT strains and the median value in cells isolated from uninfected mice. Minimum of 150 cells were evaluated for each mouse (n mice =5). **B.** Phosphorylation of histone H2AX (γH2AX) assessed by immunofluorescence analysis (green). Nuclei were counterstained with DAPI (blue). Left panel: a representative micrograph of the colonic mucosa. Scale bar: 50μM. Right panel: quantification of the γH2AX staining. At least 500 cells were evaluated for each mouse (n mice =5). The data are reported as ratio between the % of γH2AX positive cells in mice infected with the MC1Δ*cdtB* or MC1 TT strains and % of γH2AX positive cells in uninfected mice. **C.** Senescent cells (marked with black arrow heads) were detected using the SenTraGor™ reagent. Upper panel: a representative micrograph of the colonic mucosa. Scale bar: 20μM. Lower panel: quantification of the senescent positive cells. *p value ≤ 0.05, (n mice =5).

One of the consequences of DNA damage in cells exposed *in vitro* to bacterial genotoxins is the acquisition of senescence (Blazkova et al., 2010; Ibler et al., 2019), which is associated with secretion of a broad panel of mediators, mainly related to a pro-inflammatory effect, such as IL1, IL6, and IL8 (Hernandez-Segura et al., 2018). Induction of the senescent phenotype in *in vivo* bacterial infection is poorly characterized. To assess whether the DNA damaging capacity of the typhoid toxin promoted senescence in both Sv129 and C57BL6 mice, we performed hybrid histo-/immunochemical staining employing the compound GL13 (also termed SenTraGor™) that detects lipofuscin, a non-degradable aggregate of oxidized molecules that accumulates in the cytoplasm of senescent cells (Evangelou et al., 2017). As shown in Figures 2C and S3A, we detected a significant increase in GL13 epithelial positive cells in the colon and small intestine of mice infected with the MC1 TT and MC71 TT strains compared to the levels detected in mice infected with the control *Salmonella*, with the most prominent effect observed in the C57BL/6 mouse strain (cp Figure 2C and Figure S3A). In our model, induction of senescence was uncoupled from the activation of a pro-inflammatory response (Figures 1, 3 and S2).

**Figure 3.**
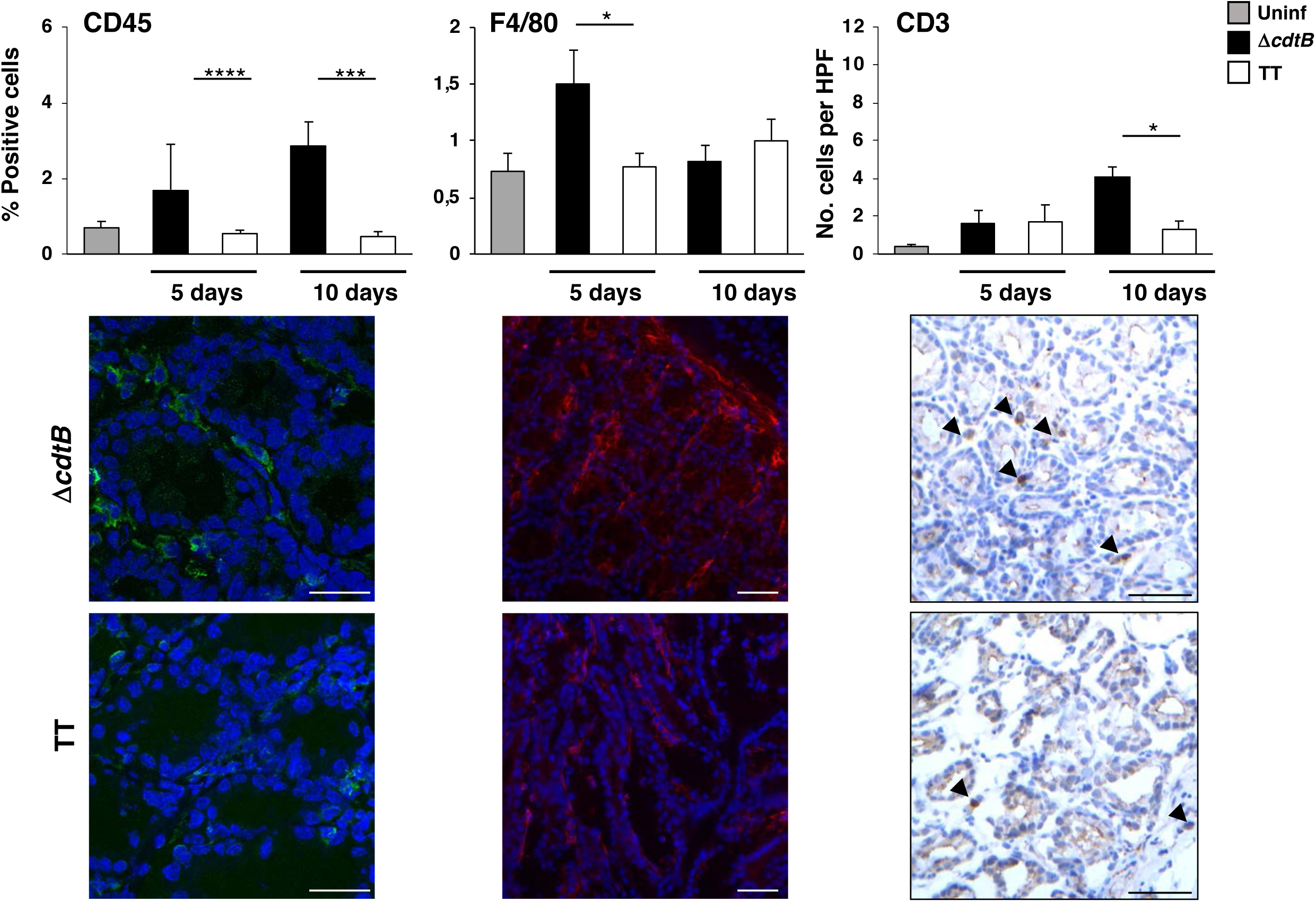
Lack of inflammatory response induced by the genotoxigenic *Salmonella* is associated with reduced levels of immune cells in the intestinal mucosa. Sv129 mice were mock infected with PBS (Uninf) or infected with the MC1 Δ*cdtB* (Δ*cdtB*) or MC1 TT (TT) strains for 5 and 10 days. Presence of immune cells was detected by immunofluorescence or histochemistry using the rabbit anti-CD45 specific antibody as leukocyte marker (left panel, green. Scale bar: 50μM); the mouse anti-F4/80 specific antibody as macrophage marker (center panel, red. Scale bar: 100μM) or a mouse anti-CD3 specific antibody as pan T lymphocyte marker (right panel, brown. Scale bar: 100μM). Upper panels representative micrographs, lower panel: quantification of the positive cells. *p value ≤ 0.05; *** p value ≤ 0.001; **** p value ≤ 0.0001, (n mice =5).

### The typhoid toxin maintains an anti-inflammatory microenvironment

Phenotypic analysis of the colonic immune cells showed a significant decrease of leukocytes (CD45) and macrophages (F4/80) already at 5 days post-infection in mice infected with the MC1 TT strain compared to the mice infected with the control *Salmonella* (Figure 3). Consistent with the time frame for activation of the adaptive immune response, infection with the MC1 Δ*cdtB* strain promoted increased levels of CD3 positive T lymphocytes at 10 days post-infection, while this effect was reduced in mice infected with the genotoxigenic strain (Figure 3).

To assess whether the genotoxigenic *Salmonella* strain suppressed the activation of the host response or promoted a qualitatively different immune response with an anti-inflammatory profile, we investigated: i) the activation of the transcription factor NFκB, by assessing the nuclear translocation of the p65 subunit (Hayden et al., 2006); ii) the presence of the non-inflammatory macrophages, by detecting mannose receptor (CD206) (Shrivastava and Shukla, 2019); iii) the levels of mRNA expression of several pro- and anti-inflammatory effector molecules by qPCR and RNAscope analysis, in the colon of uninfected controls or mice infected with MC1 TT or MC1 Δ*cdtB* strains for 5 and 10 days.

*Salmonella* infection promoted an increase in the percentage of colonic stromal cells with a nuclear pattern of p65 staining compared to the levels observed in uninfected mice, and p65 nuclear translocation was most prominent at 10 days post-infection, independently of the expression of an active genotoxin (Figure 4A and data not shown). However, in spite of a similar activation of NFκB, infection with the genotoxigenic *Salmonella* was associated with a general reduction on the mRNA levels of pro-inflammatory molecules associated with a T helper (Th) 1 response, most prominent at 10 days post-infection, and increased levels of mRNA for Th 2 cytokines associated with an anti-inflammatory environment and tissue repair (Annunziato et al., 2015; Gieseck et al., 2018; Kotas and Locksley, 2018), such as IL5, IL10 and IL13 (Figure 4B). The induction of an anti-inflammatory microenvironment was confirmed by RNAscope analysis, which showed that the levels of expression of the IL10 mRNA were higher than those of the IFNγ mRNA in C57BL/6 mice infected with the genotoxigenic *Salmonella* (Figure 4C and S3B). In accordance with the presence of a restorative response, the profile of virtually all macrophages present in the colon of mice infected with the MC1 TT strain exhibited the phenotype of non-inflammatory cells, resembling the profile observed in the colon of uninfected mice (Figure 4D).

**Figure 4.**
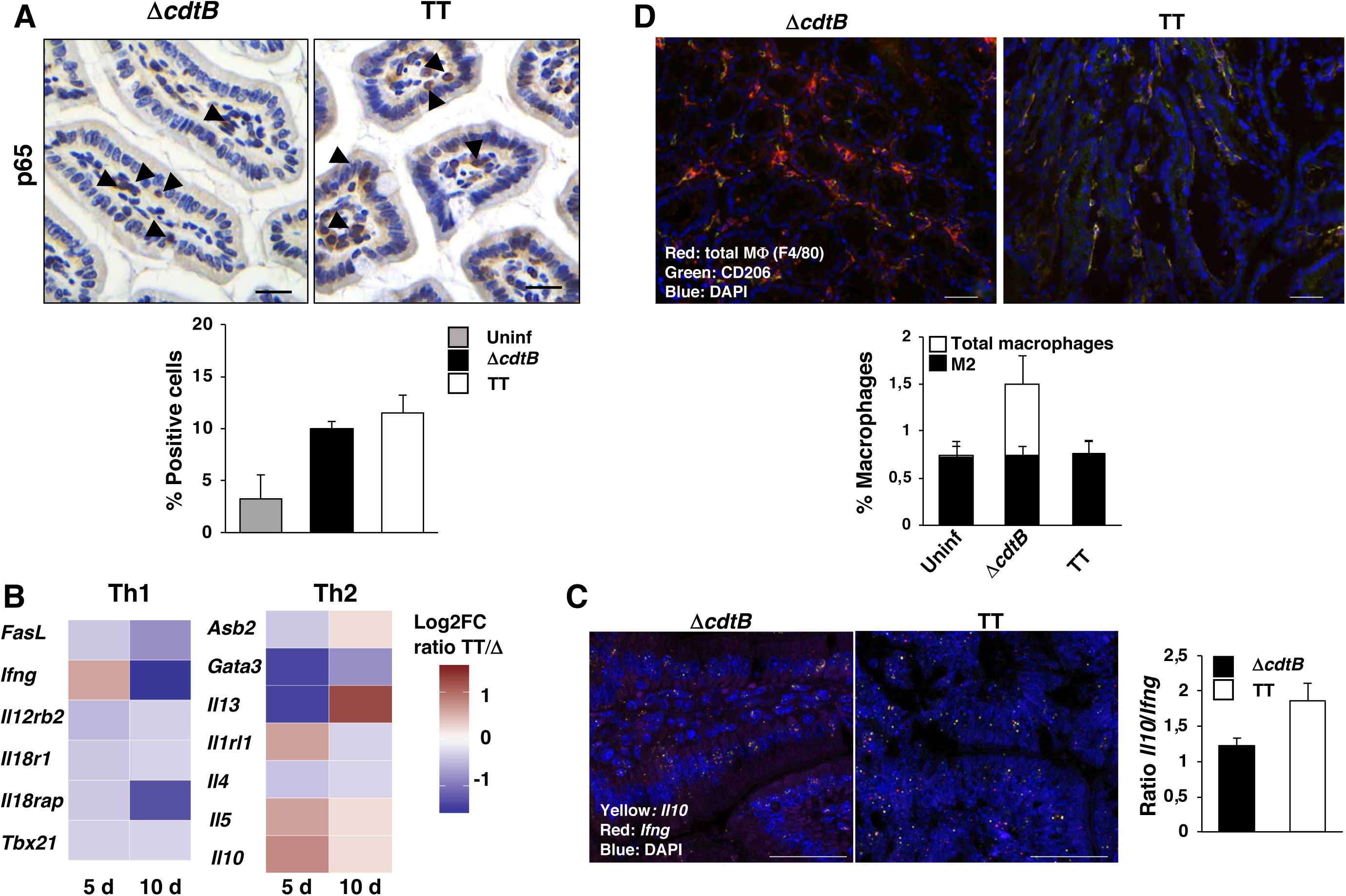
Infection with the genotoxigenic *Salmonella* induces a type 2 immune response. Sv129 mice were mock infected with PBS (Uninf) or infected with the MC1 Δ*cdtB* (Δ*cdtB*) or MC1 TT (TT) strains. **A.** Immunohistochemical analysis performed with an anti-p65 specific antibody to detect nuclear translocation of NFκB at 10 days post-infection. Upper panel: representative micrograph. Scale bar: 50μM. Lower panel quantification of cells with p65 nuclear localization (n mice =5). **B.** qPCR-Array analysis of 84 genes involved in T cell differentiation/polarization in colon of mice at 5 and 10 days post-infection. Log_2_ fold changes of mice infected with the MC1 TT strain versus the MC1 Δ*cdtB* strain (n mice =5) are displayed for Th1 and Th2 genes, respectively. **C.** RNAscope analysis to assess the levels of expression of mRNA specific for IL10 (yellow) and IFNγ (red) in C57BL6 mice infected with the MC71 Δ*cdtB* (ΔcdtB) or MC71 TT (TT) strains. Left panel representative confocal micrograph. Scale bar: 100μM. Right panel quantification of the level of mRNA expression. The data are presented as ratio between the levels of *Il10* mRNA and the levels of *Ifng* mRNA. **D.** Analysis of the macrophage population was performed with the pan-macrophage F4/80 specific antibody, followed by an TRITC-conjugated secondary antibody (red), and the CD206 antibody for non-inflammatory macrophages, followed by FITC-conjugated secondary antibody (green). Nuclei were counterstained with DAPI (blue) in the intestinal mucosa at 5 days post-infection. Left panel: representative micrograph. Scale bar: 100μM. Right panel quantification of the F4/80 and CD206 positive cells.

The immune profile data were reproduced in C57BL/6 mice infected with the MC71 TT strain, which presented a similar extend of NFκB activation associated with an enrichment of CD206 positive macrophages and regulatory T lymphocytes (T_reg_), expressing the FOXP3 transcription factor (Whibley et al., 2019) compared to the effects observed in mice infected with the control strain (Figure S3A).

We did not detect any significant difference in the percentage of cells expressing the proliferative or apoptotic markers respectively Ki67 and cleaved caspase 3 (Figure S4).

Having established a link between the presence of a functional genotoxin and a modulation of the host response, we assessed whether this effect is regulated by the ATM kinase (Shiloh and Ziv, 2013). To this end, C57BL/6 mice ATM wild type (wt) or heterozygotes for the *ATM* gene (denoted as ATM) were infected with the MC71 TT strain for 10 days. We performed immunohistochemical analysis with specific markers for activation of DDR (γH2AX), senescence (GL13), NFκB (p65), non-inflammatory macrophages (CD206) and T_reg_ (FOXP3).

No significant macroscopic difference in the histological score of the small intestine was detected (Figure S5). However, we observed a significant increase in the percentage of cells positive for γH2AX and a reduced induction of senescence (Figure 5A), indicating an inefficient activation of a proper DDR and DNA repair. This effect was associated with a reduced translocation of the NFκB subunit p65, a tendency to reduced levels of non-inflammatory macrophages and a significant reduction in the number of T_reg_ (Figure 5A), indicating the ATM kinase is required for an efficient activation of an anti-inflammatory response. Failure to activate this type of response was associated with a reduced clearance of the MC71 TT *Salmonella* strain in MLN, liver and spleen (Figure 5B).

**Figure 5.**
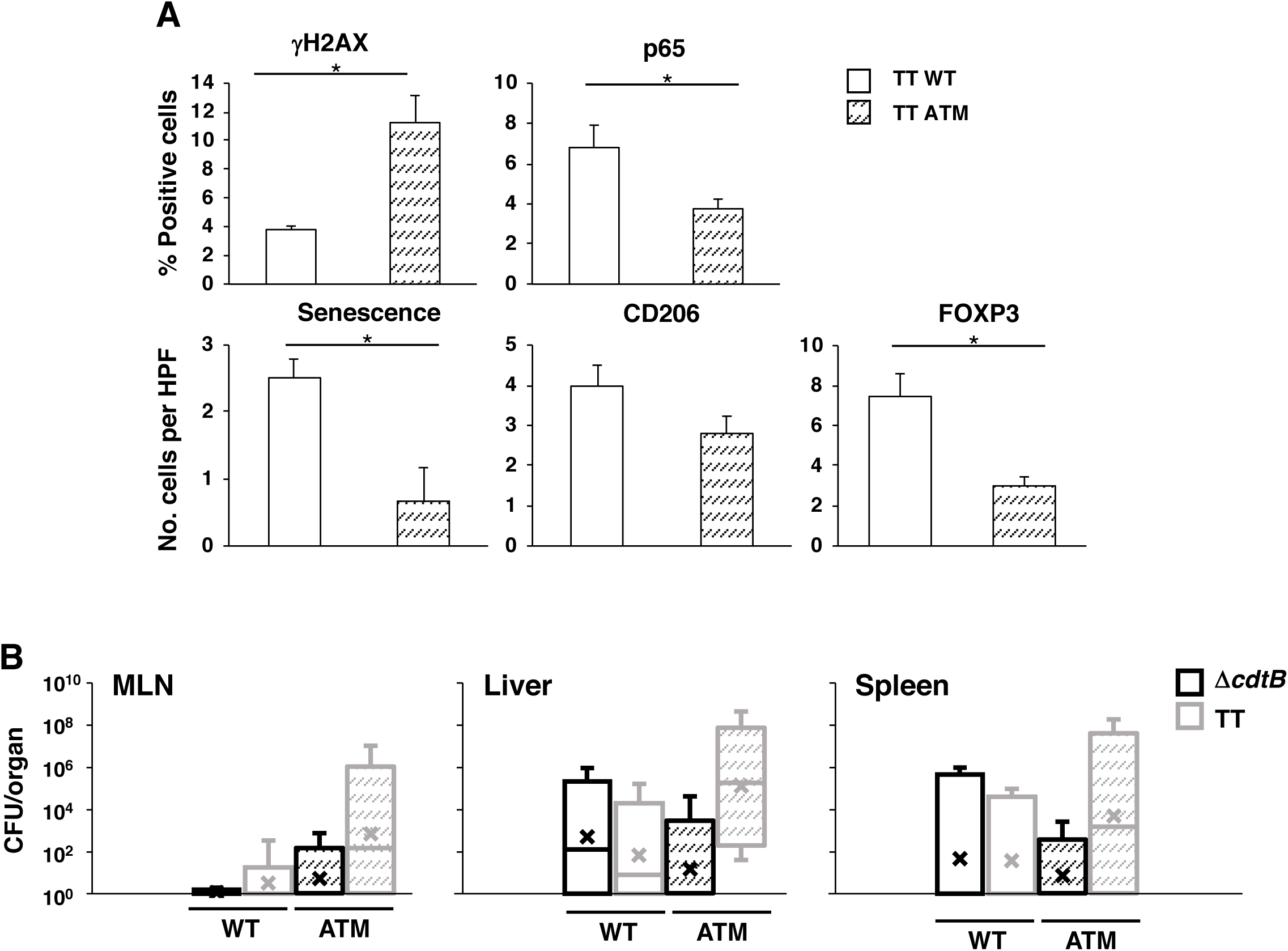
Absence of a functional ATM alters the response to the genotoxigenic *Salmonella*. C57BL/6 wild type (WT) or ATM heterozygotes (ATM) mice were infected with the MC71 Δ*cdtB* (Δ*cdtB*) or MC71 TT (TT) strains for 10 days. **A.** The following parameters were assessed by immunohistochemistry analysis: phosphorylation of H2AX (γH2AX), senescence, activation of NFκB (nuclear translocation of p65), presence of non-inflammatory macrophages (CD206) and regulatory T cells (FOXP3). **B.** Dissemination of *S.* Typhimurium in mesenteric lymph nodes (MLNs), liver, and spleen. Data are presented as colony forming unit (CFU) per organ, (n mice =5).

### The anti-inflammatory effect of the typhoid toxin is lost in subjects affected by colitis

Epidemiological evidences indicate that genotoxin producing bacteria are enriched in the microbiota of IBD and CRC patients (Buc et al., 2013; Chumduri et al., 2016; Nowrouzian and Oswald, 2012), however it is not known whether the typhoid toxin influences the course of these diseases. To address this question, we have infected Sv129 mice, suffering from Dextran Sulfate Sodium salt (DSS)-induced colitis, which is a model for a form of IBD known as Ulcerative Colitis (UC) (Chassaing et al., 2014), with the MC1 TT or the control MC1 Δ*cdtB* strains for 5 and 10 days, as summarized in Figure S6A. Infection with both *Salmonella* strains did not affect the disease activity index and amount of fecal lipocalin-2, an inflammatory marker (Chassaing et al., 2012) (Figure S6B). Histological analysis showed that a pre-existing colitis completely prevents the typhoid toxin-induced tissue protective immune response observed in healthy mice (Figure 6). The effect detected at histological levels was paralleled by the failure of the genotoxigenic bacterium to reduce the levels of leukocytes (CD45), macrophages (F4/80) and CD3 T lymphocytes in infected mice. However, we observed a time shift in the recruitment of leukocytes and macrophages, which peaked at 5 days post infection in mice infected with the control *Salmonella* strain and at 10 days post-infection in uninfected and MC1 TT infected mice (Figure 7A). We did not observe differences in the percentage of cells positive for the senescence marker GL13, suggesting that high levels of inflammation, induced by DSS, promote a senescent phenotype associated with inflammation-induced DNA damage (Wei and Ji, 2018) (Figure 7A). Interestingly, colitis enhanced significantly the colonization with the genotoxin-producing bacterium in the colon and a similar trend was also observed in the mesenteric lymph node, although the difference was not statistically significant (p=0.063) (Figure 7B).

**Figure 6.**
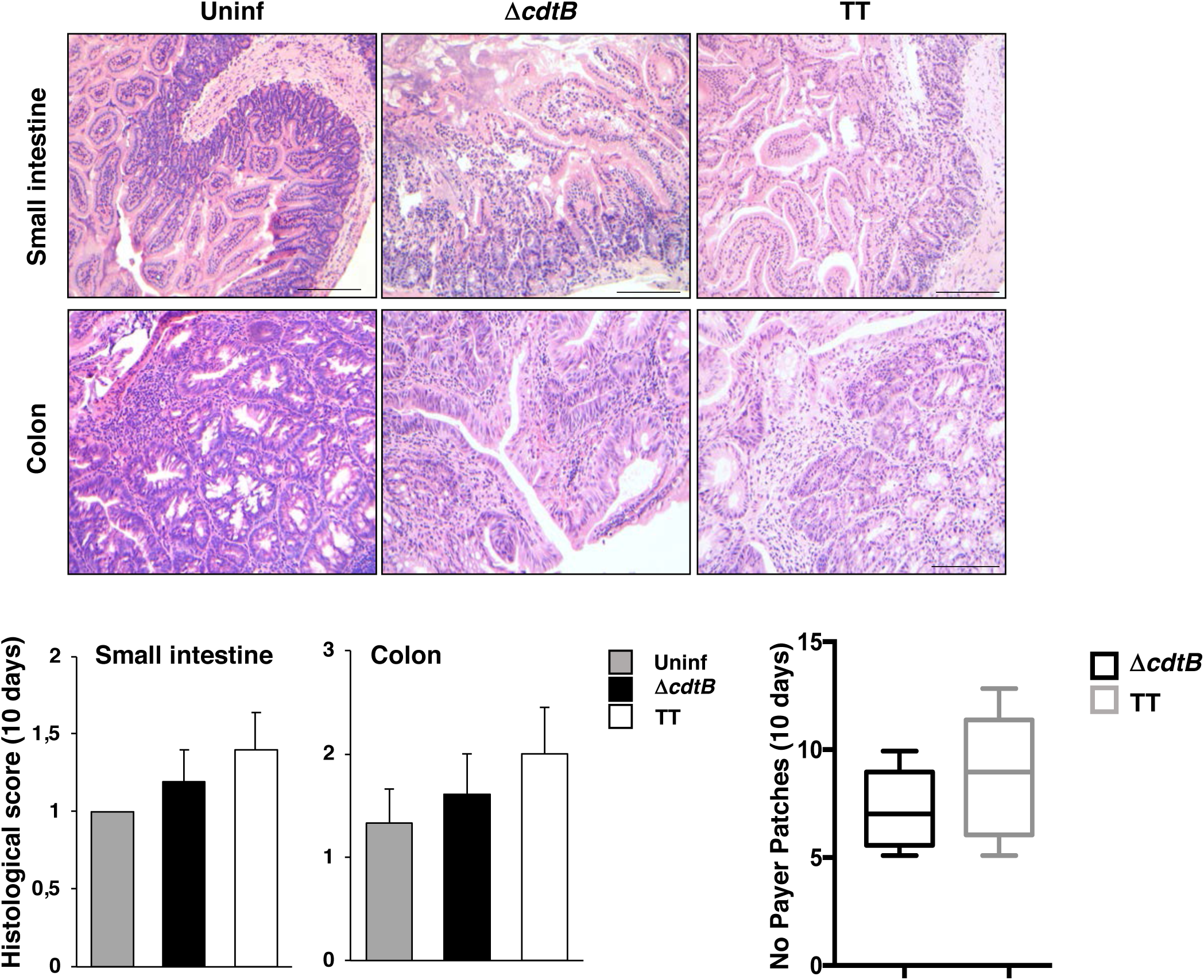
Pre-existing colitis abolishes the anti-inflammatory effect of the genotoxigenic *Salmonella*. Sv129 mice were treated with 2.5% DSS in drinking water for 7 days and then mock infected with PBS (Uninf) or infected with the MC1 Δ*cdtB* (Δ*cdtB*) or MC1 TT (TT) strains for 10 days. **A.** Haematoxylin and eosin staining of small intestine and colon. Scale bar: 200μM**. B.** Histological scores of small intestine and colon. **C.** Quantification of the macroscopically visible inflamed Peyer Patches, (n mice =5).

**Figure 7.**
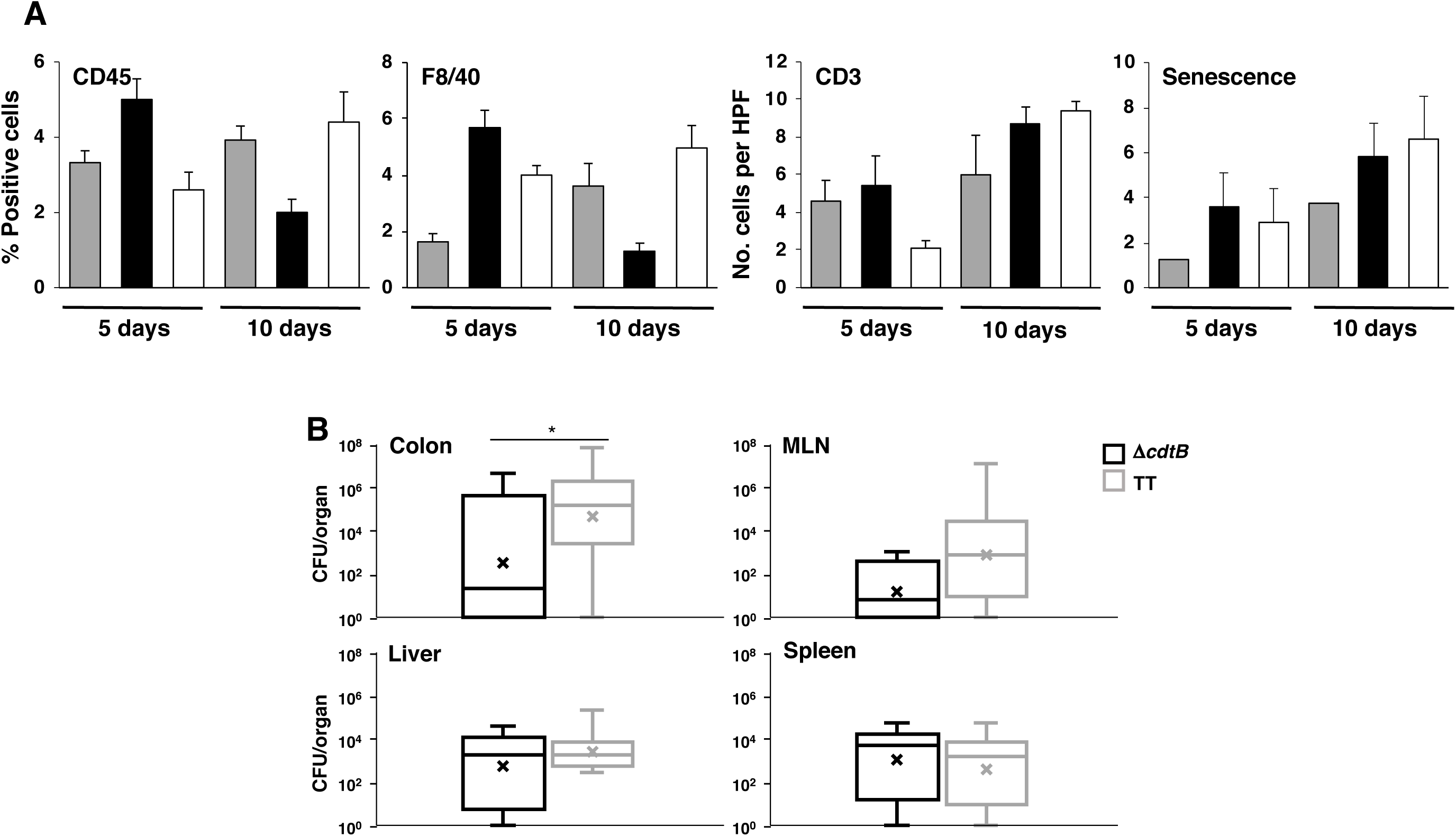
Pre-existing colitis does not prevent infiltration of immune cells but favors colonization with the genotoxic *Salmonella*. Sv129 mice were treated with 2.5% DSS in drinking water for 7 days and then mock infected with PBS (Uninf) or infected with the MC1 Δ*cdtB* (Δ*cdtB*) or MC1 TT (TT) strains for 5 and 10 days. **A.** The presence of leukocytes (CD45), macrophages (F4/80), T lymphocytes (CD3) and senescent cells was assessed as described in Figures 2 and 3, (n mice =5). **B.** Bacteria dissemination in colon, mesenteric lymph nodes (MLNs), liver, and spleen was assessed at 10 days post-infection. Data are presented as colony forming unit (CFU) per organ, (n mice =8, from two independent infection experiments).

## Discussion

We have shown that infection with *S.* Typhimurium expressing a functional typhoid toxin suppresses specifically intestinal inflammation in otherwise healthy mice (Del Bel Belluz et al., 2016). We have confirmed this observation (Figures 1 and S2), and demonstrated for the first time that a genotoxin induces DNA fragmentation in an *in vivo* model (Figure 2). Interestingly this effect was not associated with a strong induction of γH2AX, which was higher in the mucosa of mice infected with the control strain (Figure 2). These data suggest that the type of DNA damage induced in the two conditions is different: oxidative stress-induced DNA double strand breaks (DSBs) in the context of a strong inflammatory response (Kay et al., 2019), which trigger H2AX phosphorylation, and single strand breaks (SSBs) induced by low doses of the typhoid toxin, which can be converted to DSBs during S phase, as previously shown for the *E. coli* CDT (Fedor et al., 2013). It is likely that in non-inflammatory conditions the number of proliferating cells in the intestinal mucosa is low, thus limiting the number of γH2AX positive cells.

We further show that the presence of a functional toxin maintains the tolerogenic environment of the intestinal mucosa (Liu et al., 2018; Mowat, 2018), by reducing the recruitment of leukocytes, macrophages and T lymphocytes, maintaining a robust presence of non-inflammatory macrophages, T_reg_ lymphocytes, and increased mRNA levels of Th2 cytokines (Figures 3, 4, S3), in spite of a significant activation of the transcription factor NFκB and promotion of cellular senescence (Figure 2 and Figure 4). The anti-inflammatory profile (levels of T_reg_ and, to a lesser extent, macrophages) was dependent on ATM, the DNA damage transducer (Figure 5), and was lost when infection occurred in mice suffering from a pre-existing colitis (Figures 6 and 7).

These data highlight a complex interplay between the DNA damage response and the modulation of the host response upon infection with *Salmonella*, and we propose that the DDR is not only activated in a cell autonomous manner to eliminate cells with damaged DNA, but plays a non-cell autonomous role in shaping the host immune response in a tissue specific manner.

### DDR and immunomodulation

The role of the DDR and the kinases ATM and DNA-PKcs in immunomodulation is gaining a lot of interest and it has been proposed that DNA breaks and the consequent response activate a cell specific non-canonical DDR (ncDDR) that regulates the activity of these cells (reviewed in (Bednarski and Sleckman, 2019; Shiloh and Ziv, 2013)). Activation of the ATM and DNA-PKcs ncDDR in bone marrow derived macrophages is dependent on nitric oxide-induced DSB, in response to *Listeria monocytogenes* infection or stimulation with Toll-like receptor agonists (LPS). This response triggers a genetic program characterized by upregulation of a broad panel of chemokines, cytokines and inflammasome activation, with consequent production of IL1β and IL18 (Morales et al., 2017). The relevance of the ATM-dependent inflammasome activation to promote efficient bacterial clearance was shown by the enhanced susceptibility to pulmonary infection by *Streptococcus pneumoniae* in ATM deficient mice (Erttmann et al., 2016). Interestingly, activation of the ncDDR in macrophages requires signaling from the type I interferon receptors, indicating that the pathway activated by type I IFN precedes (Morales et al., 2017) and is not a consequence of the formation of DNA breaks.

The activation of an immunoregulatory ncDDR may explain why bacteria have acquired, maintained and horizontally transferred bacterial effectors that can promote DNA damage, which does not kill immediately most of its target cells *in vitro* (reviewed in (Grasso and Frisan, 2015)) and does not promote apoptosis above the levels observed in the mucosa of uninfected mice (Figure S4).

Our results support a scenario where activation of the ATM-dependent DDR promotes activation of an anti-inflammatory immune response, more prone to a tissue protection and repair (Annunziato et al., 2015; Gieseck et al., 2018; Kotas and Locksley, 2018). This scenario is supported by the observation that ATM deficiency increases the mortality rate upon DSS-induced colitis (Westbrook and Schiestl, 2010). This effect is associated with upregulation of mRNA for proinflammatory cytokines (e.g. TNFα, IL12β, IL23, IL6) and higher percentage of activated CD44 positive T cells, a marker strongly associated with the Th17 population. In line with this observation, inhibition of ATM skews the polarization of T lymphocytes derived from rheumatoid arthritis patients toward Th1 and Th17, imposing a hyperinflammatory phenotype (Yang et al., 2016). In addition, DNA damage induced by low doses of epirubicin induces an ATM-autophagy-dependent protective effect in the lung tissue of septic mice, resulting in an enhanced survival (Figueiredo et al., 2013).

Based on these observation, it is conceivable that the DNA damage response can exert multiple functions leading to different outcomes, depending on the context: i) activation of an inflammatory response triggered by the cGAS-STING or via activation of the interferon and NFκB signaling pathways by components of the DNA damage sensors such as the DNA-PK, ATM, and Mre11 (Hartlova et al., 2015; Li and Chen, 2018; Nakad and Schumacher, 2016) to eliminate dangerous cells; ii) activation of cell specific genetic programs aimed at an efficient activation of anti-microbial immunity (Erttmann et al., 2016; Morales et al., 2017); iii) fine tuning of the inflammatory response to avoid extensive inflammation-induced tissue damage (Figueiredo et al., 2013; Westbrook and Schiestl, 2010; Yang et al., 2016). Consequently, bacteria may have hijacked this immunomodulatory function and the microbial-induced DNA damage promotes stealth invasion and persistent infections (Del Bel Belluz et al., 2016). The effect observed for the typhoid toxin may also be applied to the other members of this family, where the toxin-induced DNA damage modulates the host immune response to promote a suitable niche for the microbe, and the end result of this immunomodulatory activity can vary depending on the bacterium life-style and tissue tropism.

The open question is how the ATM-induced DDR can promote different outcomes ranging from pro-inflammatory to tissue protective. It is conceivable that both the extent of the DNA damage and the immunological profile of the tissue affected by the insult exert a relevant role in determining the host response (see also section on Senescence and inflammation).

### Senescence and inflammation

Cellular exposure to genotoxins *in vitro* has been shown to induce cellular senescence (Blazkova et al., 2010; Ibler et al., 2019; Secher et al., 2013). SASP is mainly regulated by the proinflammatory transcription factor NFκB (Ohanna et al., 2011), triggered by the ATM-activation of the DDR, as well as by C/ERBPβ and MAPKp38 (reviewed in (Ohanna et al., 2011)). The secretome of senescent cells can promote an alteration of the local microenvironment ranging from maintaining tissue homeostasis (e.g. tissue remodeling or repair), generating a tumorigenic barrier to prevent cellular transformation or exerting tumor promoting effects, in a context-dependent manner (reviewed in (Faget et al., 2019; Rodier and Campisi, 2011)).

Interestingly, the composition of SASP is very complex and can differ depending on the cellular origin as well as the senescent-inducing stimulus, however the pro-inflammatory cytokines IL1 and IL6 are reported as the commonly secreted mediators in DNA damage-induced senescence (reviewed in (Gorgoulis et al., 2019)). In line with these observations, genotoxin-induced senescent cells *in vitro* secrete a plethora of mediators that support their pro-inflammatory role (Blazkova et al., 2010; Secher et al., 2013). Our data demonstrate that the presence of a functional typhoid toxin promotes senescence *in vivo*, but this effect is uncoupled from induction of a pro-inflammatory environment in healthy individuals (Figures 1, 2, S2 and S3). It is noteworthy that the anti-inflammatory effect of the MC1 TT strain was observed only in the intestine and only when infection occurs in healthy mice but not in subjects with pre-existing colitis ((Del Bel Belluz et al., 2016) and Figures 1, S2, 6 and 7). These data suggest that the type of cells (intestinal epithelial cells) and/or the type of tolerogenic environment, characterized by T_reg_ lymphocytes and non-inflammatory macrophages may impose a different type of SASP, driven by other transcription factors with or without the cooperation of NFκB. A possible candidate is the JAK2-activated STAT3 transcription factor, which has been shown to promote immune suppressive SASP profile in PTEN deficient prostate cancer, characterized by secretion of IL10 and IL13 (Toso et al., 2014), two of the Th2 cytokines whose mRNA levels were upregulated in mice infected with the MC1 TT strain (Figure 4). Considering the tissue specific and context specific response (intestine vs liver and healthy vs colitis), it is possible that the fine tune of the DDR response and the consequent SASP phenotype is modulated by the immunological profile of a specific microenvironment: a tissue-based education of senescence, which parallels the concept of tissue class control of the immune response (Matzinger and Kamala, 2011). This issue can be addressed by multiplex single cell analysis using specific markers for senescence concomitant with characterization of the cytokine profile and cell specific markers and by comparing the outcome in the intestine and liver of infected healthy mice or mice with pre-existing colitis.

### Concluding remarks

We have highlighted a complex crosstalk between the DDR induced by the typhoid toxin and the regulation of the host immune response in bacterial infection, which is not only associated with sensing dangerous signal and activating the innate immunity, but also shapes the type of immune response shifting from a pro-inflammatory response normally induced by *S.* Typhimurium toward the activation of tissue protective response. The outcome resembles infection with *S.* Typhi, characterized by lack of intestinal inflammation and a stealth invasion of the mucosa (Hiyoshi et al., 2018). However, our data highlight that the anti-inflammatory effect is context-dependent, since it is lost in subjects with pre-existing inflammatory conditions. In addition, the presence of senescent cells in absence of a strong inflammatory response in healthy subjects suggests that the overall effect of the senescence secretory phenotype depends on the tissue specific microenvironment.

Several questions still need to be addressed, including the role of the immunological profile in the definition of the SASP, the impact of chromatin alterations including γH2AX dynamics on SASP heterogeneity, the characterization of tissue and context-dependent types of senescence, and the long term consequences of infection with the genotoxigenic *Salmonella* in subjects with pre-existing colitis or ATM haploinsufficiency, where the bacterium could not be efficiently cleared (Figures 5A and 7).

## Author contributions

OCBM, ISP, TF designed the research

OCBM; DB, EP, SDPT, MMH, EB-R, AB, MLC, AW, ISP, TF performed the experiments

OCBM, DB, MMH, TT, AB, MLC, VG, TF, ISP analyzed data

TM, MR provided mouse and bacterial strains

TF wrote the manuscript with the help of OCBM and ISP

All authors commented on the manuscript

## Acknowledgements

We are grateful to Drs. Lisa Del Bell Belluz and Boris Mihaljevic for the technical assistance with the infection of the ATM mice and the RT^2^ profiler PCR array, respectively. We acknowledge the Biochemical Imaging Center Umeå (BICU) at Umeå University and the National Microscopy Infrastructure, NMI (VR-RFI 2016-00968) for providing assistance in microscopy. This investigation was supported by grants from the Swedish Cancer Society, the Swedish Research Council, the Kempestiftelserna, the Cancer Research Foundation in Northern Sweden and Umeå University to TF.

## STAR Methods

### Ethical consideration

All animals were handled in strict accordance with good animal practice as defined by the relevant national animal welfare bodies, following proceedings described in EU legislation. This study was approved by the Regional Animal Studies Ethical Committees, Northern Norrland and Stockholm Norra, Sweden (reference number A17-17 and N112/16) and from the Animal Care Committee, Toronto, Canada (reference number AUP #4088.0).

### Bacterial strains

The *Salmonella* Typhimurium strains MC1 and MC71 of the line SR-11 carrying the typhoid toxin genes and the isogenic control strains were previously described (Del Bel Belluz et al., 2016).

### DSS treatment

For the colitis-mimicking experiment, DSS 2.5% (Sigma-Aldrich Merck) was administered in drinking water *ad libitum* for 7 days prior administration of the bacterial strains or vehicle as described above. The Disease Activity Index (DAI) was determined by scoring changes in weight loss (score: 0, none; 1, 1–5%; 2, 5–10%; 3, 10–20%; and 4, > 20%), stool consistency (score: 0, normal; 2, mildly soft; 4, diarrhea), and bleeding (score: 0, normal; 2, traces of blood in stool and 4, visible rectal bleeding). All parameters were scored daily during all the experiment. The fecal lipocalin was assessed by ELISA according to the manufacturer instruction (Duoset murine Lcn-2 ELISA kit, R&D Systems, Minneapolis, MN) and as previously described (Chassaing et al., 2012).

### Infection

129S6/SvEvTac (herein named Sv129) mice were obtained from Taconic Biosciences Inc. (Bomholt, Denmark). C57BL/6 ATM deficient mice were previously described (Herzog et al., 1998). Mice were housed in a pathogen-free facility, and female mice were infected at 8 weeks of age. We used the *Salmonella* MC1 strains at the infection dose of 10^8^ for the 129S6/SvEvTac mice, and the attenuated MC71 strains at the infection dose of 10^4^ for the C57BL/6 wild type and ATM mice. Infection was performed as previously described (Del Bel Belluz et al., 2016). At 5 or 10 days post infection, intestine, liver, spleen, and mesenteric lymph nodes were collected from each mouse, as described in (Del Bel Belluz et al., 2016). For histological and immunohistochemical analysis, the tissue was either fixed in 4% paraformaldehyde and paraffin embedded (FFPE) or frozen on dry ice in a cryomold, embedded in Tissue-Tek OCT (Sakura Finetek, Tokyo, Japan).

### Histological analysis

The histopathological evaluation of intestinal inflammation was performed on hematoxylin-eosin stained sections according to previously published criteria (Erben et al., 2014). Collectively for the histopathological assessment we employed a scoring system based on the inflammatory cell infiltrate, epithelial changes and mucosal architecture (Erben et al., 2014).

### Senescence staining

Examination of senescence was performed by a hybrid histo-/immuno chemical assay utilizing GL13 (commercially available as SenTraGor®) which is a lipophilic, biotin-linked Sudan Black-B (SBB) analogue (Evangelou et al., 2017). For GL13 evaluation we counted the number of positive cells per HPF (magnification 400x).

### Comet assay

Genotoxicity on colon mucosa was evaluated by alkaline comet assay. Colon mucosa cells were collected by scraping and stored in HBSS/EDTA 0.024 M buffer at pH 7.5 before slow freezing at −80°C. Alkaline comet assay was performed as previously described (Perdry et al., 2018). Image analysis were performed with CometScore 2.0 free software. For each cell, the extent of DNA damage was calculated as percentage of pixels in the comet tail as previously described (Perdry et al., 2018). For each animal, the median of minimum 150 cells was determined.

### Immunofluorescence and immunohistochemistry

Formalin-fixed paraffin-embedded 6 μm tissue sections were deparaffinized in xylene and rehydrated through a graded series of alcohol. The endogenous peroxidase activity was blocked by incubating the sections 30 min with 3% H_2_O_2_ in distilled H_2_O.

The OCT-embedded tissues were cut with a CryoStar™ NX70 Cryostat into 6 µm thick sections at −20°C, mounted on SuperfrostTM Plus microscope slides, and stored at −80°C until usage.

#### Immunofluorescence (fresh frozen tissue)

The intestinal sections, small intestine and colon were fixed in −20°C cold acetone for 10 min. After fixation, the sections were washed two times in Tris-buffered saline (TBS) for 5 min and permeabilized with 0.1% Triton 100X, 2% fetal bovine serum (FBS), 1% bovine serum albumin (BSA) in TBS. Subsequent washing in TBS for 2 min was performed and the tissues were incubated with blocking solution (2% FBS, 1% BSA in TBS) for 30 min, followed by incubation with the primary antibodies. After incubation with the primary antibodies for 1 h, the slides were washed two times for 2 min with TBS and incubated for 1 h with the compatible secondary antibodies, used at a dilution of 1:1000 in blocking solution. Subsequently, the slides were washed again three times with TBS for 5 min and finally mounted. The nuclei were counterstained with VECTASHIELD® containing DAPI (Vector Laboratories). All images were acquired with a Zeiss Laser Scanning Microscopy LSM-510 META Confocal Microscope, equipped with a 40X Plan-Neo/1.3 NA objective. At least 500 cells were evaluated for each mouse.

#### Immunohistochemistry (FFPE)

Antigen retrieval was performed by heat-mediated antigen retrieval method in 10 mM citric acid (pH 6.0). The primary antibodies were incubated at 4°C overnight. After incubation with the primary antibodies, the slides were washed 3 times with TBS, and incubated with the appropiate HRP-conjugated secondary antibody for 1 h at a dilution of 1:200 in TBS, supplemented with 1% BSA. Immunocomplexes were visualized with the 3,3’-diaminobenzidine (DAB) substrate kit (SK-4100, Vector Laboratories Inc., Burlingame, CA, USA), according to the instructions of the manufacturer. Sections were counterstained with hematoxylin solution (Sigma-Aldrich). The evaluation of γH2AX, p65, Ki67 and cleaved caspase 3 was performed by assessing the percentage (%) of positive epithelial cells. For the assessment of CD3, FOXP3, and CD206, we counted the number of positive cells in the stroma per High Power Field (HPF, magnification 400x). Slide examination was performed by three independent observers with minimal inter-observer variability.

### Antibodies

The following primary antibodies were used for immunofluorescence analysis: rabbit anti-CD45 IgG (1:100, Abcam), rat anti-F4/80 IgG (1:100, Bio-Rad), goat anti-CD206 (1:400, R&D system), rabbit anti-γH2AX (1:200, Cell Signaling Technology).

The following primary antibodies were used for immunohistochemistry: anti-CD3 (Abcam, 1: 150), anti-CD206 (Abcam, 1:1000), anti-FOXP3 (Novus Biologicals, 1:1200), anti-p65 (Santa Cruz; 1:100), Ki-67 (Abcam, 1:100), cleaved caspase 3 (Cell Signaling, 1:100), anti-γH2AX (Ser139, Millipore; 1:1000). Antibodies were also sourced using BenchSci (www.benchsci.com).

### RT2 profiler PCR array

Total RNA from individual mice was isolated using RNeasy Mini Kit (Qiagen), pooled in equimolar amounts for each experimental group, retrotranscribed using RT^2^ First Strand Kit (Qiagen) according to the manufacturer’s instructions and loaded onto qPCR plates (Mouse Cancer Inflammation & Immunity Crosstalk and T cell subsets PCR Array, Qiagen), and as previously described (Del Bel Belluz et al., 2016).

### RNAscope

*In situ* detection of mRNA levels for IL-10 and IFNγ using the RNAscope Multiplex Fluorescent Reagent Kit (Bio-techne, Oxford, UK) was performed according to manufacturer’s protocol.

Housekeeping genes *Polymerase II Subunit A (Pol2a), Peptidylpropyl isomerase B* (*PPIB*) and *Ubiquitin C (UBC)* were used as internal controls to assess the mRNA quality. The sections were pretreated using Protease IV for 15 minutes at room temperature and further incubated with the probes targeting *Il10* and *Ifng* diluted according to manufacturer’s protocol for 2 h at 40°C.

For detection of the targeting probes, the sections were incubated with fluorescent dyes Opal 570 and Opal 690 (1:1000, #NEL810001KT, PerkinElmer). The slides were washed using the supplied washing buffer (Bio-techne, Oxford, UK) after each hybridization step at room temperature. Nuclei were counterstained with DAPI. Images were acquired with a confocal scanning microscope (Leica TCS SP8, Leica Microsystems, Wetzlar, Germany). Expression levels were assessed quantitatively using H-score, following manufacturer’s guidelines.

### Statistical analyses

All the statistical analyses have been performed using Prism 7, Graph Pad Software. The significance of differences between two experimental groups (mice infected with the control and the genotoxin *Salmonella* strain) was determined by the Student *t-*test. The significance of differences between three experimental groups was determined by ANOVA with Fisher’s LSD post-test. p values <0.05 were considered significant.

## Supplementary Information

**Supplementary Figure 1.**
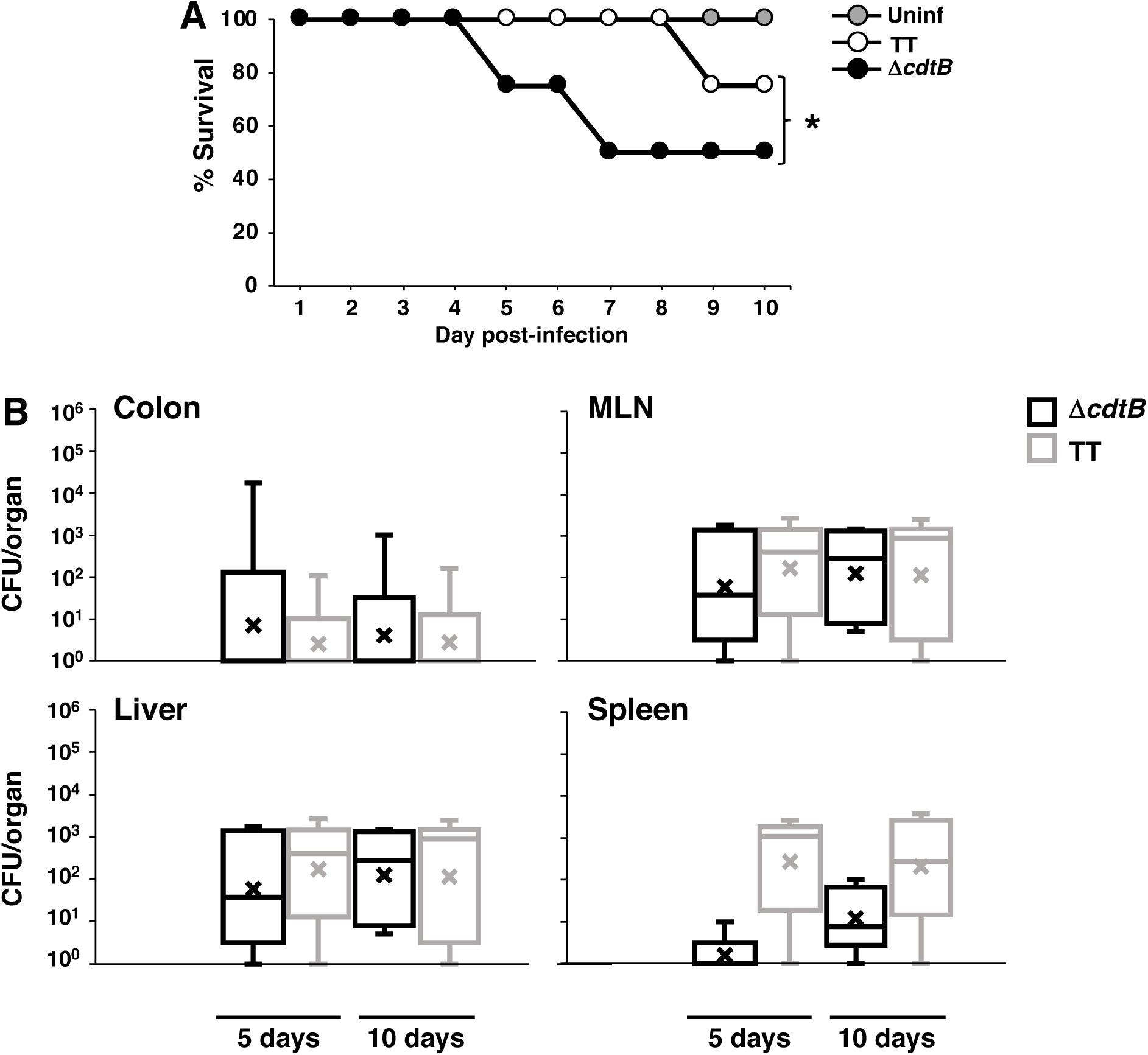
A functional typhoid toxin enhances survival rates but does not alter the extent of bacterial dissemination. **A.** Percentage of survival during the infection with the MC1 Δ*cdtB* (Δ*cdtB*) or MC1 TT (TT) strains, *p value ≤ 0.05 (n mice =5) **B.** Dissemination of the MC1 Δ*cdtB* or MC1 TT strains in colon, mesenteric lymph nodes (MLNs), liver, and spleen of Sv129 mice at 5 and 10 days post-infection. Data are presented as colony forming unit (CFU) per organ, (n mice =5).

**Supplementary Figure 2.**
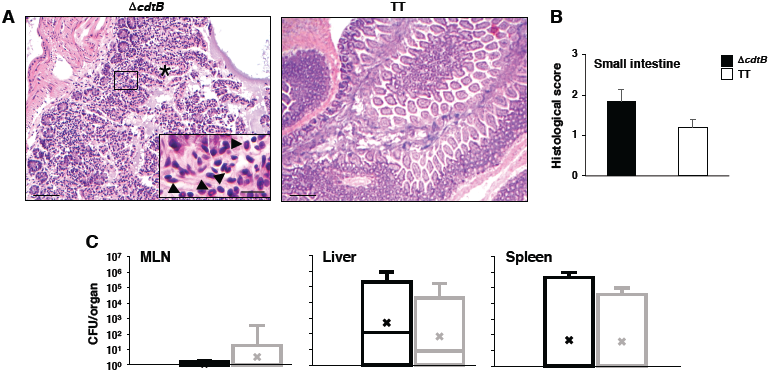
Infection with the genotoxigenic *Salmonella* prevents inflammation in C57BL/6 mice. C57BL/6 wild type mice were infected with the MC71 Δ*cdtB* (Δ*cdtB*) or MC71 TT (TT) strains for 10 days. **A.** Haematoxylin and eosin staining of the small intestine. Scale bar: 500μM (inset 50μM). **B.** Histological scores of small intestine. **C.** Dissemination of the MC71 Δ*cdtB* (Δ*cdtB*) or MC71 TT (TT) strains in mesenteric lymph nodes (MLNs), liver, and spleen at 10 days post-infection, (n mice =5).

**Supplementary Figure 3.**
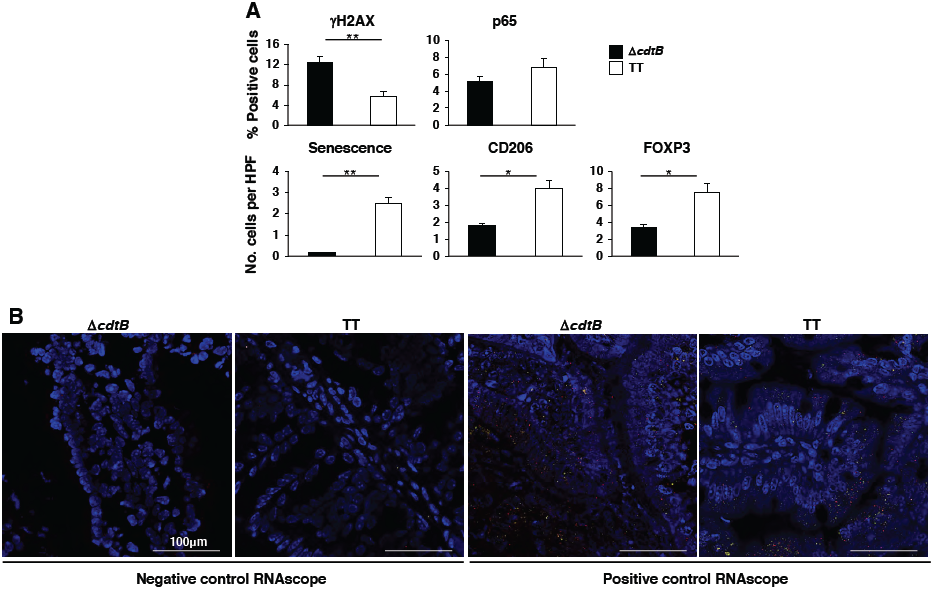
Infection with the genotoxigenic *Salmonella* induces a type 2 immune response in C57BL/6 mice. C57BL/6 wild type mice were infected with the MC71 Δ*cdtB* (Δ*cdtB*) or MC71 TT (TT) strains for 10 days. **A.** Immunohistochemistry analysis. The following parameters were assessed: phosphorylation of H2AX (γH2AX), activation of NFκB (nuclear translocation of p65), senescence, presence of non-inflammatory macrophages (CD206) and regulatory T cells (FOXP3). **B.** Negative and positive controls for assaying the probe specificity and the quality of mRNA for the RNAscope analysis. Positive controls: *Polr2a* (low expression, green), *PpiB* (intermediate expression, red), *Ubc* (high expression, yellow). Scale bar 100μM**B**

**Supplementary Figure 4.**
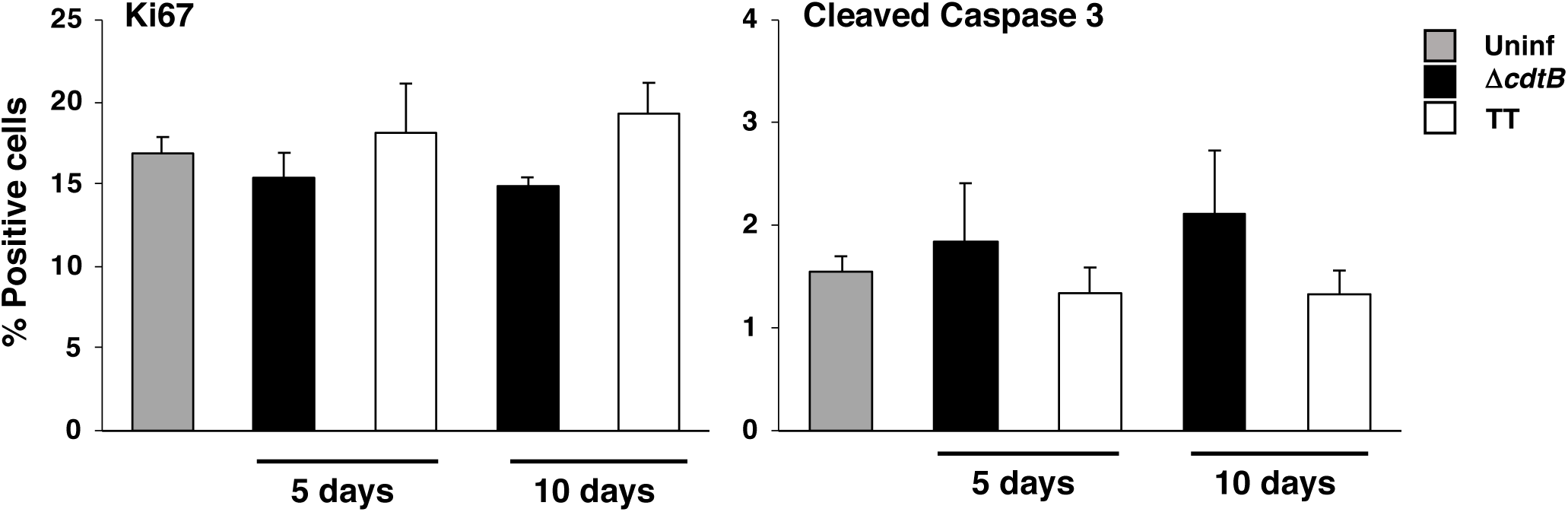
Infection with the genotoxigenic *Salmonella* does not alter the proliferative and apoptosis rate in the intestinal mucosa. Sv129 mice were mock infected with PBS (Uninf) or infected with the MC1 Δ*cdtB* (Δ*cdtB*) or MC1 TT (TT) strains for 5 and 10 days. The levels of proliferation and apoptosis were assessed by immunohistochemical analysis using the anti-Ki67 and anti-cleaved caspase 3 specific antibodies, respectively, (n mice =5).

**Supplementary Figure 5.**
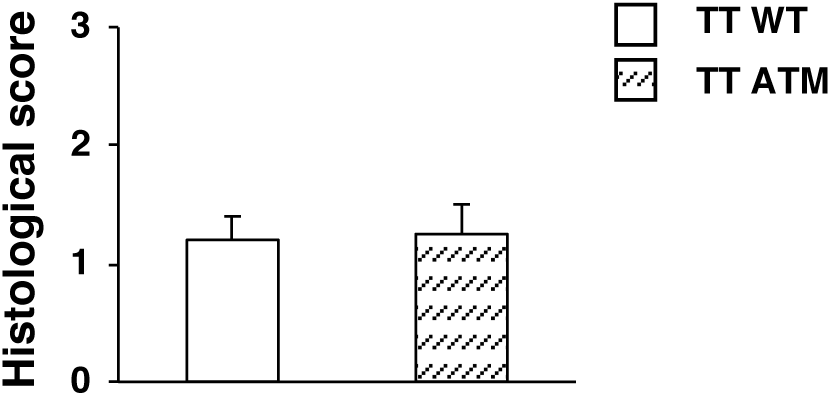
ATM deficiency does not alter the histological score in the small intestine. C57BL/6 wild type (WT) or ATM heterozygotes (ATM) mice were infected with the MC71 MC71 TT (TT) strain for 10 days and the histological score in the small intestine was evaluated.

**Supplementary Figure 6.**
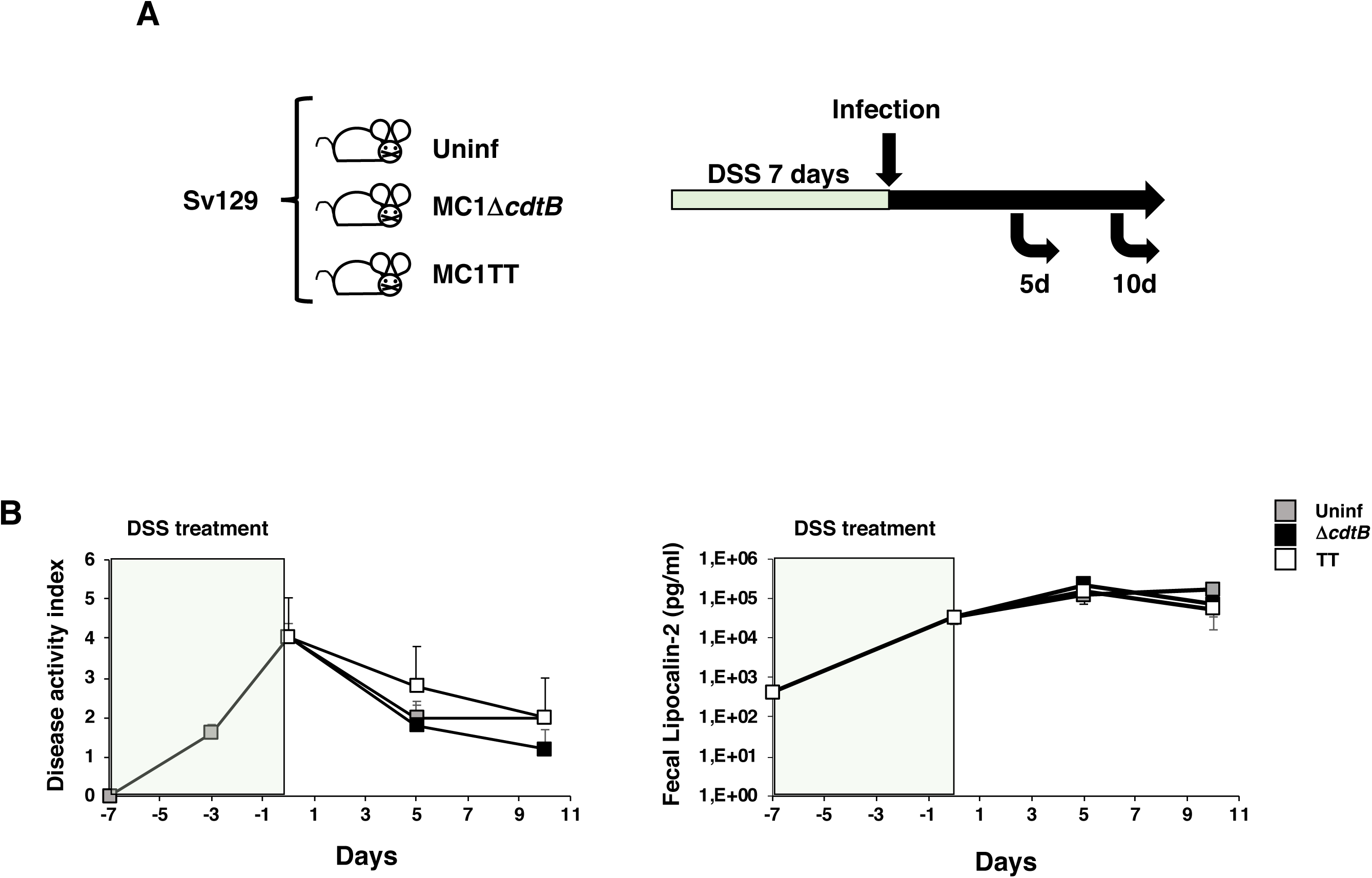
Infection with the genotoxigenic *Salmonella* does not modify the course of the DSS induced-colitis. **A.** Outline of the experimental set up. Sv129 mice were treated with 2.5% DSS in drinking water for 7 days and then mock infected with PBS (Uninf) or infected with the MC1 Δ*cdtB* (Δ*cdtB*) or MC1 TT (TT) strains for 10 days. **B.** Quantification of the disease index and the presence of Lipocalin-2 in feces, (n mice =5).

